# BMP4 regulates asymmetric Pkd2 distribution in mouse nodal immotile cilia and ciliary mechanosensing required for left–right determination

**DOI:** 10.1101/2024.05.18.594812

**Authors:** Takanobu A. Katoh, Tim Lange, Yoshiro Nakajima, Kenta Yashiro, Yasushi Okada, Hiroshi Hamada

**Affiliations:** Department of Cell Biology, Graduate School of Medicine, The University of Tokyo, Tokyo 113-0033, Japan; Laboratory for Organismal Patterning, RIKEN Center for Biosystems Dynamics Research, Kobe 650-0047, Japan; Division of Anatomy and Developmental Biology, Department of Anatomy, Kyoto Prefectural University of Medicine, Kyoto 602-8566, Japan; Laboratory for Cell Polarity Regulation, RIKEN Center for Biosystems Dynamics Research, Suita, Osaka, Japan; Department of Physics, Universal Biology Institute and International Research Center for Neurointelligence, The University of Tokyo, Hongo, Tokyo, Japan

**Keywords:** Left–right symmetry breaking, Mouse nodal immotile cilia, Pkd2, STED microscopy, Optical tweezers, Mechanobiology

## Abstract

**Background:** Mouse nodal immotile cilia mechanically sense the bending direction for left–right (L–R) determination and activate the left-side-specific signaling cascade, leading to increased *Nodal* activity. Asymmetric distribution of Pkd2, a crucial channel for L-R determination, on immotile cilia has been reported recently. However, the causal relationship between the asymmetric Pkd2 distribution and direction-dependent flow sensing is not well understood. Furthermore, the underlying molecular mechanism directing this asymmetric Pkd2 distribution remains unclear.

**Results:** The effects of several recombinant proteins and inhibitors on the Pkd2 distribution were analyzed using super-resolution microscopy. Notably, bone morphogenetic protein 4 (BMP4) affected the Pkd2 distribution. Additionally, three-dimensional manipulation of nodal immotile cilia using optical tweezers revealed that excess BMP4 caused defects in the mechanosensing ability of the cilia.

**Conclusions:** Experimental data together with model calculations suggest that BMP4 regulates the asymmetric distribution of Pkd2 in nodal immotile cilia, thereby affecting the ability of these cilia to sense the bending direction for L–R determination. This study, for the first time, provides insight into the relationship between the asymmetric protein distribution in cilia and their function.

## 1 Introduction

Establishment of left–right asymmetry in vertebrate embryos first occurs in the left–right organizer, known as the node in mice.^1^ The node is a small cavity located at the midline and is composed of two types of ciliated cells: pit cells in the central region of the node that possess motile cilia^2^ and generate the leftward nodal flow responsible for establishing left–right determination^3^ and crown cells in the peripheral region of the node that possess immotile (primary) cilia.^4^ Leftward nodal flow mechanically activates the Pkd2 channel localized on immotile cilia,^5,6^ leading to an increase in the frequency of calcium transients only on the left side of the cilia and cytoplasm.^7,8^ Particularly, leftward nodal flow causes passive mechanical bending of immotile cilia in an asymmetrical manner, wherein the left- and right-side cilia show ventral and dorsal bending, respectively.^5^ A previous experiment involving the artificial manipulation of immotile cilia in three dimensions using optical tweezers revealed that nodal immotile cilia preferentially respond to ventral bending,^5^ in the same bending direction applied to left-side cilia by leftward nodal flow. Therefore, only the left-side immotile cilia are activated by left-ward nodal flow. Activation of left-side immotile cilia induces degradation of *Dand5* mRNA, whose protein acts as an antagonist to *Nodal* and activates left-side-specific *Nodal* activity in crown cells.^9^

The Pkd2 channel was recently reported to exhibit asymmetric localization on immotile cilia, which may play a crucial role in the initial step of left–right symmetry breaking. Precise measurement of the membrane strain distribution during ventral bending induced by nodal flow suggested a relationship between the preferential response to ventral bending and the asymmetric Pkd2 distribution. Notably, passive ventral bending induces an increase in strain on the dorsal side of the cilia, where Pkd2 is preferentially localized.^5^ Therefore, the nodal immotile cilia may specifically sense dorsal bending. Dorsal localization of Pkd2 on cilia may be associated with the dorsoventral (D–V) and/or midline–lateral polarity. The asymmetric distribution of flagellar proteins, such as Hv1,^10^ has been reported in human sperm; however, the mechanism by which the asymmetric Pkd2 distribution is generated in immotile cilia remains unknown.

Pit cells, located at the center of the node, exhibit planar cell polarity along the anterior– posterior axis,^11,12^ and play a role in generating leftward nodal flow.^3,13–15^ Although cell polarity along the midline to lateral direction has not been evaluated, some factors exhibit a gradient from the midline to lateral regions. These factors may be involved in regulating the asymmetric distribution of Pkd2. For instance, Noggin and Chordin, an antagonists of bone morphogenetic protein 4 (BMP4),^16^ are expressed in the node and notochord, whereas BMP4 is expressed in the lateral plate mesoderm (LPM) from the emergence to disappearance of the node. We previously reported that exogenous BMP2 does not significantly affect Pkd2 localization on immotile cilia,^5^ however, BMP4 is expressed in the LPM. The interaction of Noggin and Chordin with BMP4 may generate a concentration gradient. Another potential candidate is the sonic hedgehog (SHH), which is expressed in the node and notochord.^17^ SHH, an extracellular protein involved in numerous signaling pathways, may generate a concentration gradient from the node to LPM. Notch2 is expressed near the node, whereas delta-like 1 is expressed in the area surrounding the node.^18^ These two genes regulate hair-bundle polarity in lateral-line hair cells.^19,20^ Hence, they may contribute to the establishment of polarity along the midline to lateral direction in crown cells. Furthermore, fibroblast growth factor 8 (FGF8) is expressed in the primitive streak of E7.5 embryos^21^ and may affect D–V polarity. We predicted that these factors establish the concentration gradient and/or polarity responsible for the asymmetric distribution of Pkd2.

Additionally, the scaffold protein in cilia may be involved in generating and maintaining the asymmetric distribution of Pkd2. Notably, the Pkd2-mastigoneme complex forms a linear array in *Chlamydomonas.*^22–24^ Although mastigonemes have not been reported in nodal cilia, Pkd2 may require other binding partners and/or scaffolds, such as Inversin. *Inversin* mutant mice exhibit *situs inversus,*^25^ and Inversin, with its compartment composed of a fibrilloid structure, may function as a scaffold.^26^ In this study, we evaluated the contribution of these factors to the asymmetric distribution of Pkd2. We also examined the relationship between the mechanosensory ability and organized distribution of Pkd2 in mouse nodal immotile cilia.

## 2 Results

### 2.1 Excess BMP4 disrupts correctly organized Pkd2 distribution in nodal immotile cilia

We examined whether the candidate factors BMP4, Noggin, Chordin, SHH signaling, Notch signaling, and FGF8 are responsible for the asymmetric distribution of Pkd2. The asymmetric distribution of Pkd2 was evaluated using confocal microscopy with an Airyscan detector after treatment with the candidates. Pkd2 in the nodal immotile cilia was observed using the *NDE2*–*hsp*– *Pkd2*–*Venus* transgene,^6^ which is driven by the nodal-specific enhancer (NDE) derived from the mouse *Nodal* gene (Figure 1A). Embryos harboring the transgene were cultured from the late bud (LB) stage, just before the beginning of node formation, to the 0–2 somite stages (ss), which is the period when symmetry breaking occurs, with or without the reagent of interest (Figure 1A). The distance along the *z*-axis between the center of Pkd2 localization and the axoneme, which represents the localization of Pkd2 protein in a cilium, was analyzed using Gaussian fitting (Figure 1B).

**Figure 1.**
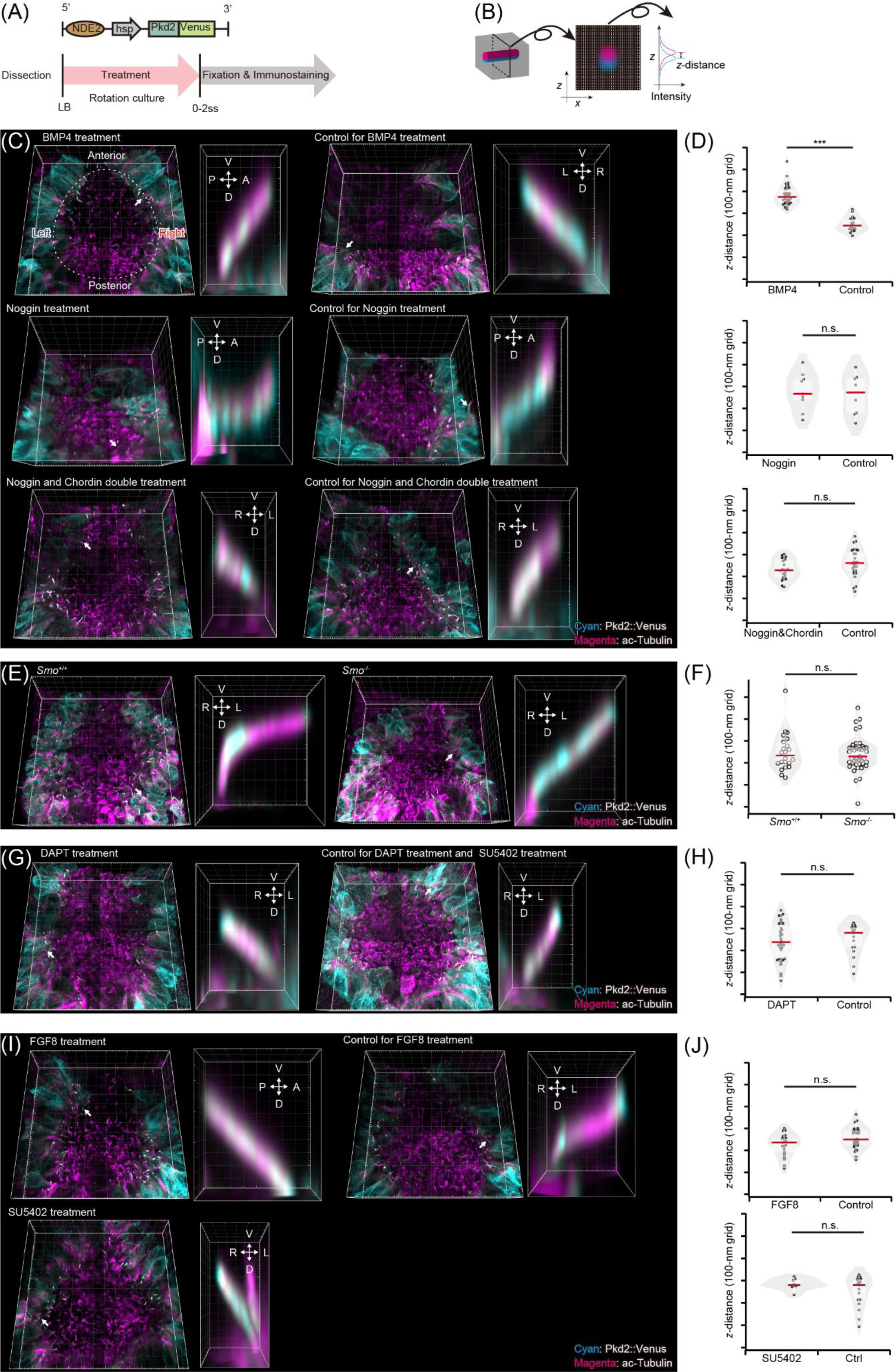
Pkd2 distribution in immotile cilia in the presence of various recombinant proteins and inhibitors and mutant embryos. **(A)** Wild-type mouse embryo harboring the *NDE2-hsp-Pkd2-Venus* transgene was cultured in agent-containing medium from the late bud (LB) stage to 0–2 somite stages (ss). The embryos were fixed with paraformaldehyde and immunostained with the Pkd2::Venus fusion protein and acetylated tubulin. Nodal immotile cilia were observed using a confocal microscope equipped with an Airyscan detector and 100× objective. **(B)** Distance between the centers of the red and green channels along the *z*-axis was measured via one-dimensional Gaussian fitting (illustrated in magenta and cyan, respectively). **(C, E, G, I)** Immunofluorescence analysis to detect the Pkd2::Venus fusion protein (cyan) and acetylated (ac) tubulin (magenta) at the node using an Airyscan microscope. Magnified images of the dorsoventral (D–V) sections of nodal immotile cilia, indicated by white arrows, are shown on the right. Grid size, 5 and 0.5 μm for the main and magnified panels, respectively. **(C)** Effect of the bone morphogenetic protein 4 (BMP4) signal on the Pkd2 distribution. Images of embryos cultured in a 1 μg/mL BMP4-containing medium (left top image), 1 μg/mL Noggin-containing medium (left middle image), 1 μg/mL Noggin- and 1 μg/mL Chordin-containing medium (left bottom image), and under control conditions (right images). **(D)** Distance along the *z*-axis between the red and green channels in BMP4-treated embryos was larger (133 ± 122 nm) than that in control embryos (*n* = 32 and 19 cilia for BMP4-treatment and control, respectively; upper panel). Distance along the *z*-axis between the red and green channels in Noggin-treated embryos was slightly smaller (15 ± 316 nm) than that in control embryos, but the difference in distribution was not significant (*n* = 26 and 12 cilia for Noggin-treatment and the control, respectively; middle panel). Distance along the *z*-axis between the red and green channels in Noggin and Chordin double-treated embryos was slightly smaller (31 ± 107 nm) than that in control embryos, but the difference was not significant (*n* = 22 and 27 cilia for Noggin and Chordin double-treatment and control, respectively; lower panel). **(E)** Effect of the sonic hedgehog (SHH) signal on Pkd2 distribution. Images of *Smo*^−/−^ (left images) and control (right images) embryos. **(F)** Distance along the *z*-axis between the red and green channels in *Smo*^−/−^ embryos was slightly smaller (9 ± 205 nm) than that in control embryos, but the difference in distribution was not significant (*n* = 43 and 22 cilia for *Smo*^−/−^ and control embryos, respectively). **(G)** Effect of the Notch signal on Pkd2 distribution. Images of embryos cultured in a 200 μM *N*-[*N*-(3, 5-difluorophenacetyl)-l-alanyl]-*S*-phenylglycine *t*-butyl ester (DAPT)-containing medium (left images) and under control conditions (right images). **(H)** Distance along the *z*-axis between the red and green channels in DAPT-treated embryos was slightly smaller (11 ± 170 nm) than that in the control embryos, but the difference in distribution was not significant (*n* = 36 and 17 cilia for DAPT-treatment and control, respectively). **(I)** Effect of the fibroblast growth factor (FGF) signal on the Pkd2 distribution. Images of embryos cultured in a 1 μg/mL FGF-8b-containing medium (left upper images), 25 μM SU5402-containing medium (left upper images), and under control conditions (right images). **(J)** Distance along the *z*-axis between the red and green channels in FGF-8b-treated embryos was slightly smaller (32 ± 40 nm) than that in the control embryos, but the difference in distribution was not significant (*n* = 21 and 30 cilia for FGF-8b treatment and control, respectively). Distance along the *z*-axis between the red and green channels in SU5402-treated embryos was larger (70 ± 147 nm) than that in the control embryos (same data as the control for DAPT-treatment), but the difference in distribution was not significant (*n* = 17 cilia for SU5402 treatment). **(H, J)** Note that the control for DAPT treatment and SU5402 treatment is the same dataset. **(D, F, H, J)** Red bars indicate the median values. \*\*\**P* < 0.001; n.s., not significant (Mann– Whitney *U* test).

First, we examined the effect of BMP4 signaling by supplementing the medium with recombinant BMP4 or its antagonists, Noggin and Chordin. In embryos treated with 1 µg/mL recombinant BMP4, the distance between the center of cilia and location of Pkd2 in immotile cilia was apparently more polarized (133 ± 122 nm; mean ± standard deviation) toward the dorsal side compared with that in the control (Figure 1C). In contrast, Pkd2 localization did not significantly differ between the Noggin-treated and control embryos or between the Noggin and Chordin double-treated and control embryos (Figure 1D). We next evaluated the effects of SHH signaling in mutant embryos lacking *Smo,*^27^ which encodes a G protein-coupled receptor that plays a crucial role in the Hedgehog signaling pathway. Pkd2 localization did not significantly differ between *Smo*^−/−^ mutant and control embryos (Figure 1E, F). Finally, we investigated the involvement of Notch signaling by supplementing the medium with *N*-[*N*-(3, 5-difluorophenacetyl)-l-alanyl]-*S*-phenylglycine *t*-butyl ester (DAPT), a γ-secretase inhibitor. No significant difference was observed in Pkd2 localization between DAPT-treated and control embryos (Figure 1G, H). The effect of FGF signaling was examined by supplementing the medium with recombinant FGF8 or SU5402 (inhibitor of the tyrosine kinase activity of FGF and vascular endothelial growth factor receptors). These treatments did not significantly affect Pkd2 localization (Figure 1I, J).

The results, derived from Airyscan images, suggest that an excessive amount of BMP4 influences the proper distribution of Pkd2 in immotile cilia, with no involvement of other signaling factors. To confirm the effects of BMP4, Pkd2 localization in immotile cilia was examined using 3D-stimulated emission depletion (STED) microscopy, which achieved a resolution of <100 nm in the *x, y,* and *z* directions.^28^ We determined the angular distribution of Pkd2::Venus protein on the transverse plane of the axoneme by measuring the Venus intensity and angle from the center of gravity of acetylated tubulin (Figure 2A).^29^ Analysis of the STED images revealed that cilia in BMP4-treated embryos exhibited a significantly biased Pkd2 distribution toward the dorsal side with a D/(D + V) ratio of 0.65 ± 0.14 (*n* = 40; Figure 2B, C) compared with that in the control embryos with a ratio of 0.54 ± 0.12, as previously reported.^5^ In contrast, cilia on Noggin-treated embryos showed an asymmetric distribution with a D/(D + V) ratio of 0.55 ± 0.09 (*n* = 36), which was similar to that in the control (Figure 2B, D).

**Figure 2.**
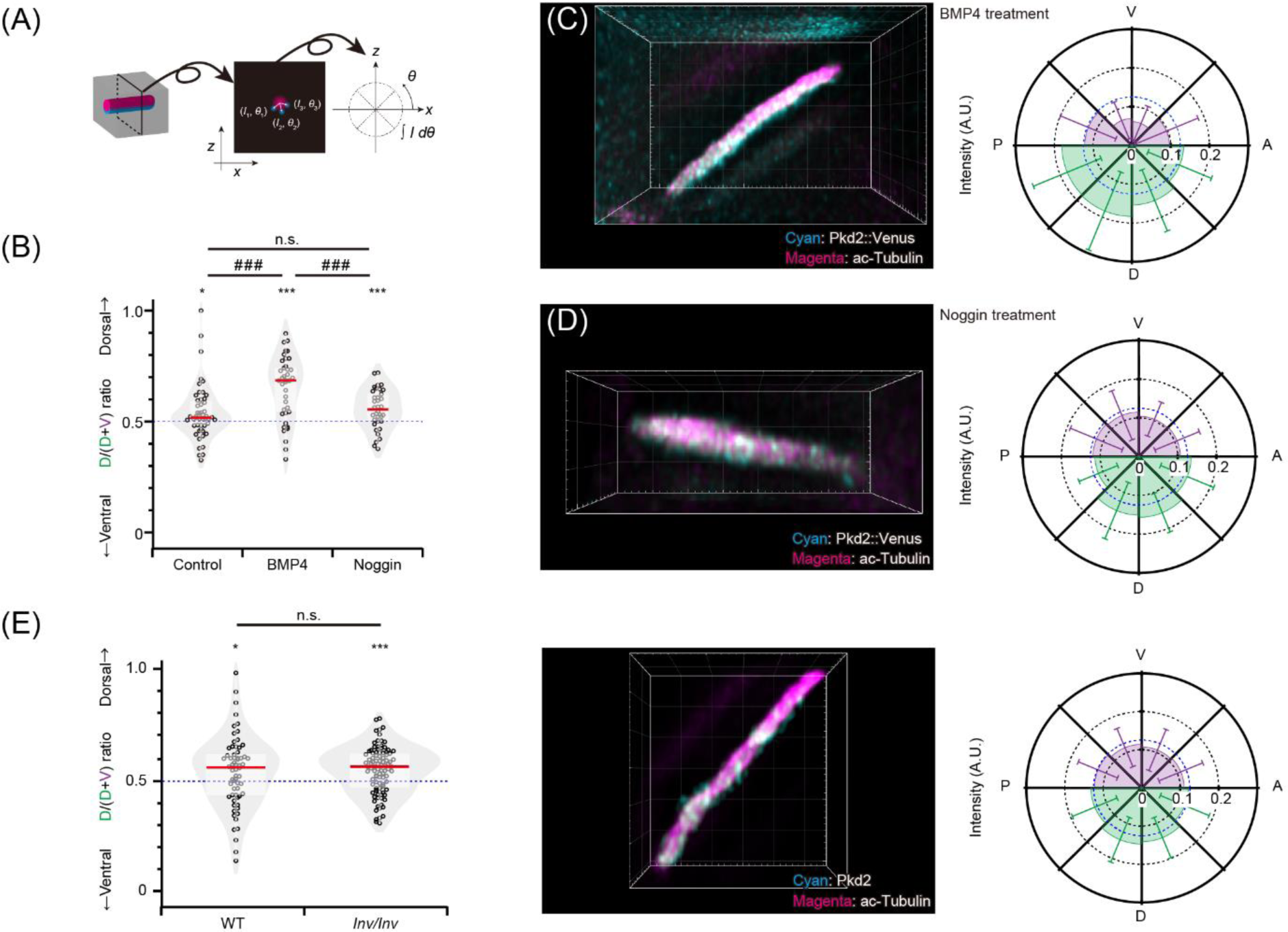
Angular distribution of Pkd2 in the presence of recombinant BMP4 or Noggin protein evaluated using 3D-stimulated emission depletion (STED) microscope. **(A)** Wild-type mouse embryos harboring the *NDE2-hsp-Pkd2-Venus* transgene were cultured in agent-containing medium from LB to 0–2 ss. The embryos were immunostained with anti-green fluorescent protein (green channel; illustrated in cyan) and anti-acetylated tubulin (red channel; illustrated in magenta) and observed using 3D-STED microscopy. Green fluorescence intensity in transverse planes of each cilium was analyzed and plotted with each 45° sector based on the gravity center of red fluorescence intensity. **(B)** Ratio of Pkd2::Venus signal intensity on the dorsal side to that on the dorsal plus ventral sides (D / [D + V]). Ratio in the cilia of BMP4-treated embryos was significantly larger than that in the cilia of previously reported wild-type embryos^5^ and Noggin-treated embryos. Interestingly, the ratio in the cilia of Noggin-treated embryos did not significantly differ compared with that of previously reported wild-type embryos.^5^ Red bars indicate the median values. \**P* < 0.05; \*\*\**P* < 0.001 (one-sample *t* test). ###*P* < 0.001; n.s., not significant (Mann–Whitney *U* test). **(C, D)** Immunofluorescence analysis to detect the Pkd2::Venus fusion protein (cyan) and acetylated (ac) tubulin (magenta) at the node using a 3D-STED microscope after BMP4 (C) or Noggin (D) treatment. Grid size, 500 nm. **(C)** Magnified views of D–V sections of BMP4-treated cilia observed using the 3D-STED microscope (left). Angular distribution of green fluorescence intensity in the transverse planes of BMP4-treated cilia was analyzed (*n* = 40 cilia; right). **(D)** Magnified views of D–V sections of Noggin-treated cilium observed using the 3D-STED microscope (left). Angular distribution of green fluorescence intensity in the transverse planes of Noggin-treated cilia was analyzed (*n* = 36 cilia; right). **(E)** Ratio of Pkd2 signal intensity on the dorsal side to that on the dorsal plus ventral sides (D / [D + V]). Ratio of the signal intensity in the cilia of *Inv/Inv* embryos was comparable to that of previously reported wild-type embryos.^5^ Red bars indicate the median values. \**P* < 0.05; \*\*\**P* < 0.001 (one-sample *t* test). ###*P* < 0.001; n.s., not significant (Mann–Whitney *U* test). Grid size, 500 nm.

To assess the association between the scaffold protein Inversin in cilia and asymmetric distribution of Pkd2, we used *Inv/Inv* mutant mice. The localization of Pkd2 in immotile cilia was evaluated using STED microscopy, employing the Pkd2 antibody rather than the *Pkd2*–*Venus* transgene. Cilia on *Inv/Inv* embryos exhibited an asymmetric distribution with a D/(D + V) ratio of 0.57 ± 0.10 (*n* = 64), which was similar to that in control embryos with a ratio of 0.54 ± 0.14, as previously reported^5^ (Figure 2E). These results indicate that excess BMP4 disrupts the correct distribution pattern of Pkd2, inducing a more polarized distribution toward the dorsal side of nodal immotile cilia.

### 2.2 Model calculation supports that the BMP4 concentration gradient is involved in an asymmetric Pkd2 distribution

We conducted model calculations to determine the effect of BMP4 on Pkd2. Initially, we estimated the concentration gradient of BMP4 in the node of both wild-type (WT) embryos and BMP4-treated embryos (Figure 3A, B). In the WT embryo model, BMP4 was secreted from the left LPM. We set the BMP4 concentration of the LPM as 1 nM; the molecules diffused with a diffusion constant of 3 µm^2^/s (Table 1). Nodal immotile cilia were located 120 µm away from the LPM. Despite considering internalization and non-specific degradation, measured as a clearance rate of 8.9 × 10^−5^/s^30^ (Table 1), some secreted BMP4 reached the node and generated a concentration gradient (Figure 3C). The estimated BMP4 concentration gradient at the nodal immotile cilia was approximately 0.001 nM/µm in WT (Figure 3D). In contrast, in the model under BMP4-treated conditions, the node depth was 12 µm, with the immotile cilia 6 µm from the bottom of the node. Considering the BMP4 clearance on the node surface (Table 1), a BMP4 concentration gradient was expected to from along the node depth (Figure 3E). The estimated BMP4 concentration gradient at the nodal immotile cilia was approximately 5.2 nM/µm under the BMP4-treated condition (Figure 3F). Based on the parameters derived from measurements in previous reports, the BMP4 concentration gradient at the nodal immotile cilia was significantly increased under the BMP4-treated condition.

**Figure 3.**
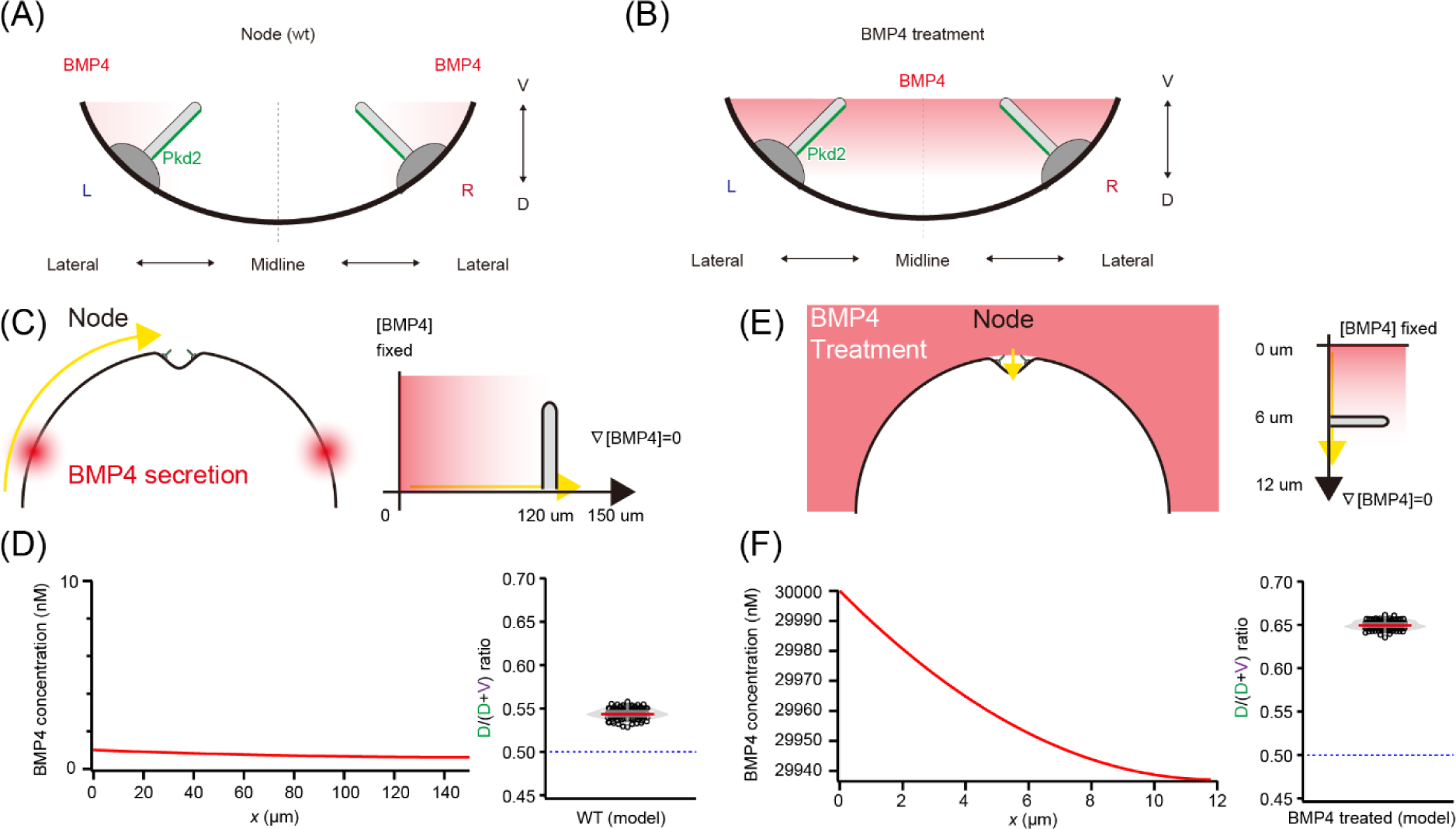
Calculation of BMP4 concentration and model of regulation of asymmetric Pkd2 distribution. **(A)** BMP4 is expressed in the lateral plate mesoderm (LPM). Endogenous expression of BMP4 generates a concentration gradient from the LPM to the node. The asymmetric distribution may be explained by this BMP4 gradient, which enables the arrangement of Pkd2 molecules away from the high BMP4 concentration regions. **(B)** In the presence of excess BMP4, the BMP4 gradient may be increased locoregionally, causing more asymmetric Pkd2 distribution and enrichment in the dorsal region. **(C)** Schematic of the model of the wild-type (WT) embryo. BMP4 was secreted from the LPM (depicted as a red sphere), with the left side of the LPM defined as *x* = 0 µm. The center of the node was positioned 150 µm from the left side of LPM, and immotile cilia on the left side were 120 µm from the left side of LPM. BMP4 concentration was maintained at 1 nM on the left side of LPM, and BMP4 diffused from this point. **(D)** Calculated BMP4 concentration along the left side of LPM to the node in the WT embryo (left). The concentration at 5000 s, after reaching a plateau, is shown. The resulting Pkd2 distribution in cilia is shown as the D/(D + V) ratio (right). **(E)** Schematic of the model of BMP4 treatment. The medium contained a high concentration of BMP4 (depicted in red). Upper side of the node was defined as *x* = 0 µm, and the node depth was defined as 12 µm. Immotile cilia were 6 µm from the bottom of the node. BMP4 concentration in the medium was maintained at 30 µM, and BMP4 diffused into the node. **(F)** Calculated BMP4 concentration from the upper to the bottom of the node under BMP4 treatment (left). The concentration at 3600 s, after reaching a plateau, is shown. The resulting Pkd2 distribution in cilia is shown as the D/(D + V) ratio (right).

**Table 1.** Model parameters (see Figure 4)

|  |  | Wild-type | BMP4 treatment |
| --- | --- | --- | --- |
| Parameters<br>for<br>calculating<br>BMP4<br>diffusion | Diffusion constant of BMP4 ($D_b$;<br>use value for BMP2b<sup>30</sup>) | 3 $\mu\text{m}^2/\text{s}$ | 3 $\mu\text{m}^2/\text{s}$ |
|  | Initial concentration of BMP4 at the<br>center of the node | 0 M | 0 M |
|  | BMP4 concentration in LPM | 1 nM | - |
| | BMP4 concentration in the medium | - | 30 $\mu\text{M}$ (1 $\mu\text{g}/\text{mL}$) |
| | Degradation constant of BMP4 ($\lambda$;<br>use value for BMP2b<sup>30</sup>) | $8.9 \times 10^{-5} \text{ /s}$ | $8.9 \times 10^{-5} \text{ /s}$ |
|  | Duration | 5000 s | 3600 s |
| Parameters<br>for modeling<br>the<br>asymmetric<br>Pkd2<br>distribution<br>in cilia | Diffusion constant of Pkd2 in cilia<br>(value for PTCH1<sup>51</sup>) | 0.1 $\mu\text{m}^2/\text{s}$ | 0.1 $\mu\text{m}^2/\text{s}$ |
| | Sensitivity of BMP4 concentration<br>gradient for Pkd2 distribution<br>regulation ($S$) | $8 \times 10^{10}$ | $6 \times 10^7$ |
| | Estimated $D/(D + V)$ ratio of Pkd2<br>(see Figure 4D,F) | $0.54 \pm 0.01$ | $0.65 \pm 0.01$ |

The relationship between the BMP4 concentration gradient and regulation of the Pkd2 asymmetric distribution remains unknown. A plausible hypothesis is that the BMP4 concentration influences the Pkd2 distribution by biasing the diffusion of Pkd2 in response to the BMP4 concentration gradient. In this hypothetical model, when the BMP4 concentration decreased toward the center (bottom) of the node, Pkd2 molecules exhibited biased diffusion, showing preferential movement toward the dorsal side. We chose ‘*S*’ (sensitivity of BMP4 concentration gradient for regulation of Pkd2 distribution; Table 1) as a model parameter to match the experimental values (see Methods). Through this parameter adjustment, the model successfully reproduced the Pkd2 asymmetric distribution observed in the experimental data (Figure 3D, F). These models indicate that the BMP4 concentration gradient influences the generation of the asymmetric Pkd2 distribution (Figure 3A, B).

### 2.3 Excess BMP4 disrupts the calcium response in crown cells triggered by mechanical stimuli to the cilium

To examine the relationship between the asymmetric Pkd2 distribution and its ability to sense the bending direction of immotile cilia, we used optical tweezers to manipulate individual immotile cilia and simultaneously measure the calcium response in the cytoplasm of crown cells.^5^ Briefly, *iv/iv* embryos harboring the *NDE4*–*hsp*–*5HT_6_*–*GCaMP6*–*2A*–*5HT_6_*–*mCherry*^5,7^ and *NDE4*–*hsp*–*GCaMP6*^5^ transgenes were cultured from the LB to 0–2 ss in the presence of recombinant BMP4 or Noggin protein (Figure 4A). The frequency of calcium transients in the cytoplasm was assessed while applying dorsal and ventral bending of the cilium using a 3.5-μm-diameter bead trapped using optical tweezers^5,29,31^ (Figure 4A). Immotile cilia in the left–right organizer of mouse^5^ and zebrafish^32^ embryos function as mechanosensors. Furthermore, in mouse embryos, these immotile cilia preferentially sense ventral bending.^5^ We postulated that the asymmetric distribution of Pkd2 is associated with this directional sensing of bending.

**Figure 4.**
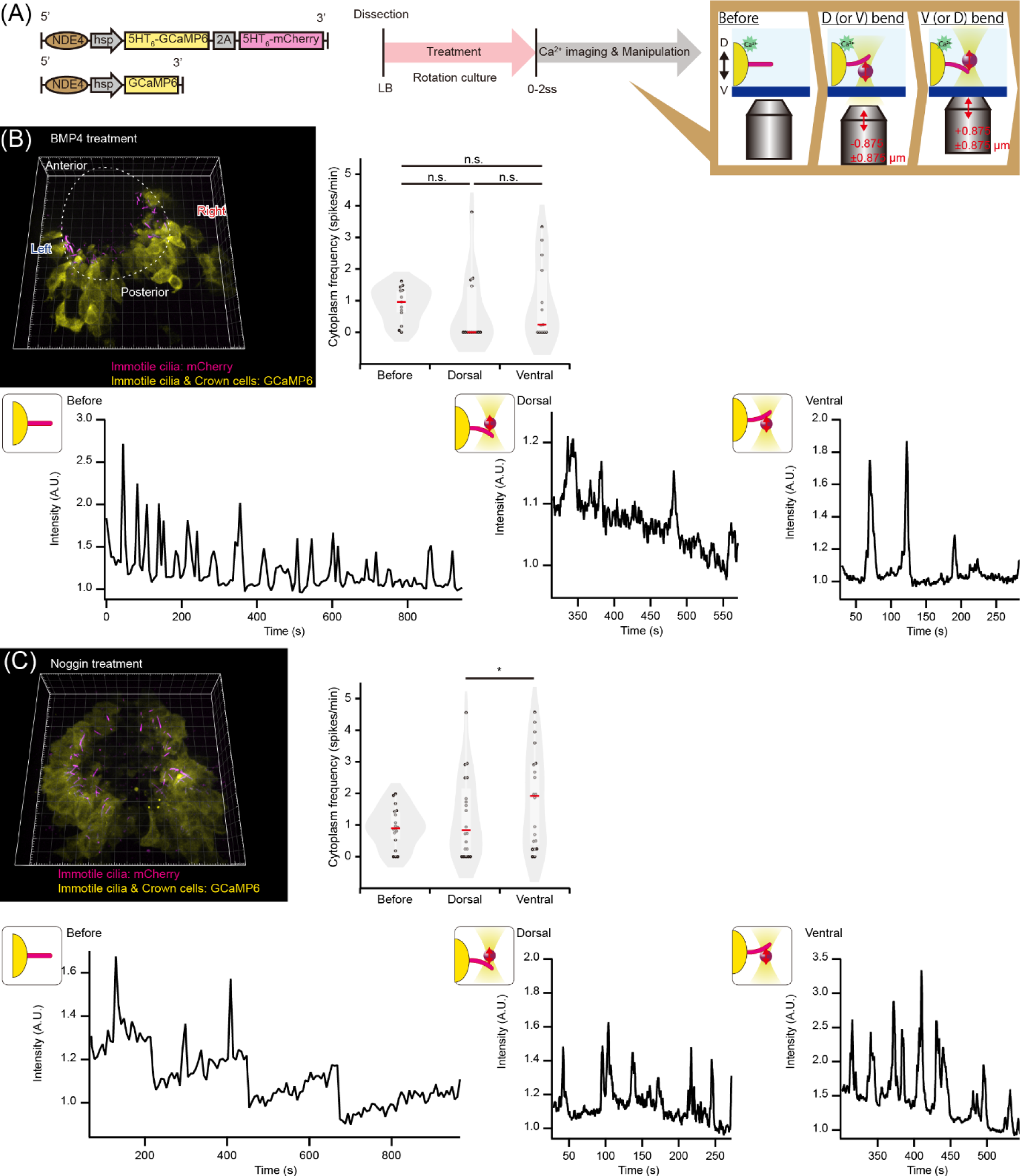
Calcium response in crown cells triggered by mechanical stimuli to the cilium in the presence of recombinant BMP4 or Noggin protein. **(A)** Experiments were performed using *iv/iv* mouse embryos harboring two transgenes. Immotile cilia at the node were visualized based on mCherry expression, which is regulated by a nodal-specific enhancer (NDE) and targeted to cilia by a 5-hydroxytryptamine receptor isoform 6 (5HT_6_) sequence (upper left). GCaMP6 was expressed in the cytoplasm for cytoplasmic calcium imaging (lower left). The embryos were cultured in agent-containing medium from LB to 0–2 ss and subjected to calcium imaging (middle panel). The cilium was subjected to sequential dorsal and ventral bending by a polystyrene bead trapped using optical tweezers. A bead was trapped, displaced to *z* = –1.75 μm (equivalent to the radius of the bead; the minus sign indicates the ventral direction according to the use of an inverted microscope), and made to contact the ventral side of the cilium. The cilium was subjected to dorsal bending of –0.875 μm at 2 Hz with an amplitude of ±0.875 μm for approximately 250 s (cilium was bent 0 to 1.75 μm toward the dorsal side; middle in the right panel) and then bent ventrally at +0.875 μm at 2 Hz with an amplitude of ±0.875 μm for approximately 250 s (right in the right panel). **(B)** Three-dimensional image of the node of a BMP4-treated embryo (left top). Grid size, 10 µm. The mean frequency of the calcium transient in the cytoplasm before stimulation, during dorsal bending, and during ventral bending (*n* = 13 cilia; right top). Average ± standard deviation (SD) values of the mean calcium frequency were 0.89 ± 0.53, 0.66 ± 1.13, and 0.98 ± 1.19 Hz before stimulation, during dorsal bending, and during ventral bending, respectively. Red bars indicate the median values. Time course of cytoplasmic calcium signal intensity in the crown cell before stimulation (left bottom), during dorsal bending (middle bottom), and during ventral bending (right bottom) (GCaMP6 *F/F*_0_ ratiometric values). **(C)** Three-dimensional image of the node of Noggin-treated embryos (left top). Grid size, 10 µm. The mean frequency of calcium transient in the cytoplasm before stimulation, during dorsal bending, and during ventral bending (*n* = 18, 20, and 20 cilia for before stimulation, during dorsal bending, and during ventral bending, respectively; right top). Average ± SD values of the mean calcium frequency were 0.93 ± 0.61, 1.27 ± 1.25, and 1.83 ± 1.49 Hz before stimulation, during dorsal bending, and during ventral bending, respectively. Red bars indicate the median values. Time course of cytoplasmic calcium signal intensity in the crown cells before stimulation (left bottom), during dorsal bending (middle bottom), and during ventral bending (right bottom; GCaMP6 *F/F*_0_ ratiometric values).

In BMP4-treated embryos, crown cells exhibited intrinsic calcium oscillations, as previously reported^5,7,8^; however, the calcium frequency under dorsal or ventral bending was not significantly increased compared with that before applying mechanical stimuli (Figure 4B). This result indicates that nodal immotile cilia did not respond to mechanical stimuli under treatment with excess BMP4. In contrast, in Noggin-treated embryos, the calcium frequency was significantly increased under ventral bending compared with that under dorsal bending or before applying mechanical stimuli (Figure 4C), similarly to in the WT embryos, as previously reported.^5^ Therefore, Noggin-treated cilia retained the ability to sense the bending direction, whereas BMP4-treated cilia lost their mechanosensing ability. Hence, correctly organized asymmetric distribution of Pkd2 in cilia may be crucial for sensing the bending direction as well as for mechanosensing ability.

## 3 Discussion

BMP4 functions in left–right determination within the context of Nodal expression in the LPM. When left-sided-specific Nodal activation occurs at the node, the signal is transmitted to the LPM.^33^ In the LPM, a self-enhancement and lateral-inhibition system involving Nodal and Lefty maintains robust Nodal expression.^34^ As BMP signaling represses Nodal expression in the LPM, the antagonism of BMP by Noggin and Chordin in the left LPM relieves the repressive effects of BMP on Nodal expression. Therefore, the asymmetric BMP signal distribution is involved in maintaining left–right asymmetric Nodal expression in the LPM.^16^ In this study, we observed an additional role for BMP4 in left–right determination. Notably, a specific concentration gradient of BMP4 is necessary to generate a correctly organized asymmetric distribution of Pkd2 on cilia, thus affecting their mechanosensing ability. The precise mechanism underlying regulation of the Pkd2 distribution by the BMP4 concentration remains unknown; however, cilia or the apical surface of crown cells may sense the BMP4 concentration gradient along the D–V axis, which would place Pkd2 molecules away from the higher BMP4 concentration (Figure 3A, B), bringing more Pkd2 molecules to the dorsal side. In contrast, Inversin, the scaffold protein in cilia, is not involved in generating the asymmetric Pkd2 distribution.

The model calculation explained why Pkd2 distribution was more polarized toward the dorsal side in the presence of excess BMP4. However, the biophysical mechanism underlying detection of the BMP4 concentration gradient, particularly sensing of the shallow gradient of 0.001 nM/µm in the WT embryos, remains unknown. Based on studies of the physics of chemoreception in microorganisms,^35^ sensing through the cell surface or a multicellular system coupled with sufficient observation (integral) time is a more plausible mechanism than a simple receptor-based sensing system within cilia. Notably, node emergence and ciliogenesis occur during the mid-streak to late streak stages.^36^ In contrast, BMP4 expression begins in the extra-embryonic mesoderm before node emergence and later detected in the posterior primitive streak and LPM at embryonic day 7.5^16,37,38^. Based on these reports, the BMP4 concentration gradient may emerge prior to node formation. Subsequently, during ciliogenesis and by utilizing this BMP4 concentration gradient, Pkd2 may gradually be asymmetrically distributed over time.

The molecular pathway responsible for regulating the Pkd2 distribution via the BMP4 concentration gradient requires further analysis. Particularly, the reasons why Noggin and Chordin did not affect the Pkd2 distribution remain unclear. Dand5, which is expressed in the crown cells and functions as an antagonist of BMP4, may lead to these effects. However, to establish a robust concentration gradient along the midline–lateral direction, BMP4 should antagonize a molecule expressed in the center of the node, such as those found in pit cells. Another possibility is the compensatory effect of BMPs, as BMPs exhibit considerable overlap in receptor-binding specificity. For instance, both BMP4 and BMP6 bind to BMP receptor-II.^39^ Notably, Noggin showed a stronger inhibitory effect on BMP4 than on BMP6 in receptor binding^40^. Following Noggin treatment, other BMPs, such as BMP6, may compensate for the function of BMP4. BMP6 is expressed in the yolk sac endoderm and anterior cardiogenic mesoderm on embryonic day 7.0 in mouse embryos,^41^ suggesting that is plays a role in generating concentration gradients from the lateral side of the embryo to the node. Pkd2 is trafficked to the cilia through its C-terminus, where it interacts with PKD1. Notably, Pkd1 harbors a cilia-targeting sequence at its C-terminus^42^. However, the precise mechanism by which the BMP4 concentration gradient regulates the Pkd2 distribution remains unclear. Certain molecules capable of sensing BMP4 may be involved in regulating the asymmetric Pkd2 distribution. To identify proteins that may interact with BMP4, we conducted a search using BioGRID,^43^ focusing on ciliary proteins,^44–46^ and identified the following nine candidates: CACL, MAP4, RPL4, RPL7, RPL9, RPL12, RPL18, RPL21, and SYNCRIP. SUFU was excluded from the list because *Smo* was not involved in regulating the Pkd2 distribution (Figure 1F). These nine candidates may be involved in regulating the Pkd2 distribution. Furthermore, BMP receptors are localized at the base of osteoblast cilia.^47^ Although the expression of these candidates in nodal immotile cilia has not been evaluated, they may be involved in regulating the asymmetric distribution of Pkd2.

Our data demonstrate that an organized asymmetric Pkd2 distribution is necessary for the mechanosensing ability of nodal immotile cilia. However, the exact mechanism of bending-direction sensing using dorsally localized Pkd2 remains unknown. We induced excessive dorsal localization of Pkd2 via BMP4 treatment. We hypothesized that the cilia exhibit an exaggerated response to ventral bending, based on our previous study, as excessive dorsal localization of Pkd2 may be more sensitive to the increase in membrane tension on the dorsal side of the cilia caused by ventral bending. However, we found that the cilia lost their mechanosensing ability rather than exhibiting an exaggerated response to ventral bending. The reason for this loss of mechanosensing ability in cilia under BMP4 treatment requires further analysis. However, this may be attributed to the loss of the mechanosensing function of Pkd2. Pkd2 functions by forming a heterocomplex with Pkd1l1 in the nodal immotile cilia.^48,49^ Excessively polarized localization of Pkd2 may prevent the correct formation of a complex with Pkd1l1 in the cilia. Considering the report on the Pkd2–mastigoneme complex in *Chlamydomonas,*^22–24^ Pkd2 may require other binding partners and/or scaffolds, such as Inversin,^25,26^ for its mechanosensing function, although Inversin was not involved in generating the asymmetric Pkd2 distribution (Figure 2E). Notably, excessive BMP4 may affect these interactions. Another possibility is that excessive BMP4 may disrupt signal transduction from the cilia to the cytoplasm. We previously reported that calcium transients in cilia trigger calcium release from the endoplasmic reticulum, which is abundant in the apical region of crown cells, in an IP3 receptor-dependent manner.^7^ This cascade then activates *Dand5* mRNA degradation, ultimately establishing left–right asymmetry^7,9^. Although reports on the effect of BMP4 on this cascade remain lacking, BMP4 may disrupt signal transduction through this pathway. Further studies are needed to understand the molecular mechanism underlying the establishment of the organized polarized Pkd2 distribution based on the BMP4 concentration gradient.

## 4 Experimental Procedures

### 4.1 Mice and transgenes

*Smo*^+/−^ mice (strain B6(D2)-Smo<RGSC2073>) were obtained from the Riken Bioresource Center (Ibaraki, Japan). Transgenic mouse lines harboring *NDE2–hsp–Pkd2–Venus*^6^, *NDE4–hsp–GCaMP6– pA*^5^, and *NDE4–hsp–5HT_6_–GCaMP6–2A–5HT_6_–mCherry*^7^ were previously reported. *Inv/Inv* mutant mice were previously reported^25^. All animal experiments were approved by the Institutional Animal Care and Use Committee of the RIKEN Kobe Branch and Committee for Animal Research of Kyoto Prefectural University of Medicine.

### 4.2 Embryo culture

Mouse embryos were collected on embryonic day 7.5 in HEPES-buffered Dulbecco’s modified Eagle’s medium (DMEM, pH 7.2). Embryos at the LB stage were selected and cultured using the roller culture method at 5% CO_2_ and 37 °C in 50-mL tubes containing DMEM supplemented with 75% rat serum. Agents were added to the medium at final concentrations of 1 μg/mL for BMP4 treatment (314-BP; R&D Systems, Minneapolis, MN, USA), 1% v/v of 4 mM HCl containing 0.1% bovine serum albumin (BSA; A3311; Sigma-Aldrich, St. Louis, MO, USA) as the control of BMP4 treatment, 1 μg/mL for Noggin treatment (1967-NG; R&D Systems), 1% v/v of phosphate-buffered saline (PBS) containing 0.1% BSA as the control of Noggin treatment, 1 μg/mL Noggin and 1 μg/mL Chordin as double treatment (758-CN; R&D Systems), 2% v/v of PBS containing 0.1% BSA as the control Noggin and Chordin double treatment, 200 µM for DAPT (AG-CR1-0016; AdipoGen, San Diego, CA, USA), 25 µM for SU5402 (CS-0200; ChemScene, Monmouth Junction, NJ, USA), 0.4325% dimethyl sulfoxide (276855; Sigma-Aldrich) as the control of DAPT treatment or SU5402 treatment, and 1 μg/mL for FGF-8b (423-F8; R&D Systems), 0.5% v/v of PBS containing 0.1% BSA as the control of FGF-8b treatment.

### 4.3 Analysis of Pkd2 localization in immotile cilia

Pkd2 localization in immotile cilia was analyzed as previously described.^5,29^ After roller culture, the embryos were washed with PBS. Immunostaining was performed as previously described.^5,29^ Anti-acetylated tubulin (1:200 dilution; T6793; Sigma-Aldrich) and anti-green fluorescent protein (1:200 dilution; ab13970; Abcam, Cambridge, UK) primary antibodies were used. The pair of secondary antibodies used were anti-chick and anti-mouse conjugated either with AlexaFluor Plus 488 (1:200 dilution; A32931; Invitrogen, Carlsbad, CA, USA) and AlexaFluor Plus 555 (1:200 dilution; A32727; Invitrogen) for Airyscan observation or STAR ORANGE (1:200 dilution; anti-chick; Abberior, Göttingen, Germany) and STAR RED (1:200 dilution; anti-mouse; Abberior) for STED observation.

The nodes were observed using a Zeiss LSM 880 (Oberkochen, Germany) with an Airyscan confocal microscope equipped with 100× (alpha-Plan-APOCHROMAT 100 Oil for SR 1.46 N.A.) lenses. For accurate measurement, the treatment and control embryos were alternately measured with in the same experiment set. 3D images were recorded with 2 × 2 tiling and depth along the *z*-axis of 168 nm. After Airyscan processing, the distance along the *z*-axis between the centers of the red and green channels was measured via Gaussian fitting with a fitting error of σ < 20 nm, as previously described^.5^ 3D-STED imaging was performed using an Infinity instrument (Abberior) equipped with a 60× lens (UPLXAPO60XO 1.4 N.A.; Olympus, Tokyo, Japan) as previously described.^5,29^ Image processing and analysis of the angular distribution of the intensity of Pkd2 signals were conducted as previously described.^5,29^ For *Inv/Inv* embryos, we used a Pkd2 antibody (1:50 dilution; sc-28331, Santa Cruz Biotechnology, Dallas, TX, USA) for immunostaining, as previously reported.^5^

### 4.4 Manipulation of cilia using optical tweezers and analysis of calcium transients

A distal portion of each embryo including the node after roller culture with agents was excised, placed in a chamber consisting of a glass slide fitted with a thick silicone rubber spacer (thickness: 400 μm), covered with a coverslip (No.1S; Matsunami), and incubated at 5% CO_2_ and 37 °C in FluoroBrite DMEM (A1896701; Thermo Fisher Scientific, Waltham, MA, USA) supplemented with 75% rat serum and agents. The experiment using optical tweezers^50^ was performed using an IX83/CSU-W1 microscope equipped with a 60× lens (UPLSAPO 60XW 1.2 N.A.; Olympus), single-mode fiber laser (wavelength of 1064 nm; YLR-5-1064-LP-SF; IPG Photonics, Novi, MI, USA), and filter set (ZT1064rdc-sp-UF3, Chroma Technology, Bellows Falls, VT, USA and SIX870; Asahi, Tokyo, Japan) as previously described.^5,29^

### 4.5 Model and numerical simulations

Diffusion of BMP4 was calculated using the following equation:

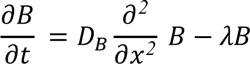

where *B*, *λ*, and *D*_*B*_ represent the concentration, degradation constant, and diffusion constant of BMP4, respectively. As values for *λ* and *D*_*B*_, we used reported values obtained for BMP2b in zebrafish embryos.^30^ Although the precise concentration of *B* in the LPM of WT embryos was unknown, it was estimated to be approximately the order of 1 nM. The values of each parameter are listed in Table 1. The geometric configurations of the models are illustrated in Figure 3C for WT embryos and Figure 3E for BMP4-treated embryos. In the model, the concentration of *B* at *x* = *0* was fixed at 1 nM for the WT and 30 µM (1 µg/mL) for the BMP4-treated condition. The initial concentration of BMP4 was 0, except where at *x* = *0*. The boundary conditions were defined by the no-flux condition at the center of the node (L) for the WT and at the bottom of the node (L) for the BMP4-treated condition: *∇B*|_*x*=*L*_ = *0*. For numerical simulations of the model, we used the Crank– Nicolson scheme with 751 grids for WT and 61 grids for BMP4-treated condition (grid width, 200 nm) and a constant time step of *Δt* = 0.001s.

The distribution of Pkd2 molecules in the cilium was modeled by simulating the diffusion of individual Pkd2 molecules along the *x*-axis, representing the dorsoventral (transverse) axis of the cilium, with the diffusion constant *D_pkd_*. The cilium was divided into 10 steps (width of each step, 0.1, corresponding to 20 nm) along the *x*-axis, and each Pkd2 molecule was moved with the following probabilities:

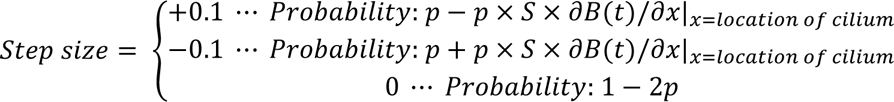

where *p* = *Δt* × *D*_*pkd*_/(*step size*)^2^ and *S* is the sensitivity of BMP4 concentration gradient for Pkd2 distribution regulation. When the BMP4 concentration gradient (*δB*(*t*)/*δx*) was 0, indicating the absence of a BMP4 concentration gradient, Pkd2 molecules exhibited free diffusion. In contrast, when *δB*(*t*)/*δx* was a negative value, indicating a decrease in the BMP4 concentration toward the center (bottom) of the node, Pkd2 molecules showed biased diffusion toward the dorsal side depending on the parameter, *S*. The diffusion constant of Pkd2 within cilia was approximated using the value of PTCH1.^51^ The parameter *S* was selected to ensure that the model replicated the observed asymmetric Pkd2 distribution observed via STED microscopy. The boundary conditions were defined by the reflective conditions at both the dorsal and ventral edges of the cilium. For numerical simulations, 100 Pkd2 molecules were initially located at the center of the cilia and then diffused according to the model, and the positions of these molecules during 4500–5000 s (for WT; Figure 3D) and 3100–3600 s (for the BMP4-treated condition; Figure 3F) were plotted as the D/(D + V) ratio.

### 4.6 Statistical analysis

Statistical analysis and graph preparation were performed using IgorPro 8 (WaveMetrics, Portland, OR, USA). Three-dimensional reconstructed images were generated using Imaris software (Oxford Instruments, Oxford, UK). All statistical tests are described in the figure legends. *P* < 0.05 was considered statistically significant.

## Acknowledgments

We would like to thank X. Sai, S. Hiver, Y. Ikawa, and H. Nishimura for providing assistance with the mouse embryo experiments, H.Y. Liu and S. Orii for help with STED image analysis; and T. Furuya for secretarial assistance. Additionally, we would like to thank K. Kawaguchi and members of his laboratory, as well as the members of the Okada laboratory, for their helpful discussion.

## Grant Sponsor

This study was supported by the FOREST Program (grant no. JPMJFR224N) of the Japan Science and Technology Agency (JST), Grant-in-Aid (grant no. 21K15096, 24K18107, and 24H01270) from the Japan Society for the Promotion of Science (JSPS), RIKEN Special Postdoctoral Researcher Program, The University of Tokyo Excellent Young Researcher Program, Ultrastructure Research Fund from The Japanese Society of Microscopy to T.A.K.; by a grant from the Japan Society for the Promotion of Science (no. 17H01435) to H.H.; by JSPS grants 19H05794, 19H05795 and 22H04926; JST grants JPMJMS2025-14, JPMJCR20E2 to Y.O. This work was supported in part by the University of Tokyo Research Internship Program (UTRIP) 2023 to Y.O.

## Conflict of Interest Disclosure

The authors declare no conflicts of interest.

## Notes

### Competing Interest Statement

The authors have declared no competing interest.

